# Improving functional protein generation via foundation model-derived latent space likelihood optimization

**DOI:** 10.1101/2025.01.07.631724

**Authors:** Changge Guan, Fangping Wan, Marcelo D. T. Torres, Cesar de la Fuente-Nunez

## Abstract

A variety of deep generative models have been adopted to perform *de novo* functional protein generation. Compared to 3D protein design, sequence-based generation methods, which aim to generate amino acid sequences with desired functions, remain a major approach for functional protein generation due to the abundance and quality of protein sequence data, as well as the relatively low modeling complexity for training. Although these models are typically trained to match protein sequences from the training data, exact matching of every amino acid is not always essential. Certain amino acid changes (e.g., mismatches, insertions, and deletions) may not necessarily lead to functional changes. This suggests that maximizing the training data likelihood beyond the amino acid sequence space could yield better generative models. Pre-trained protein large language models (PLMs) like ESM2 can encode protein sequences into a latent space, potentially serving as functional validators. We propose training functional protein sequence generative models by simultaneously optimizing the likelihood of training data in both the amino acid sequence space and the latent space derived from a PLM. This training scheme can also be viewed as a knowledge distillation approach that dynamically re-weights samples during training. We applied our method to train GPT-like models (i.e., autoregressive transformers) for antimicrobial peptide (AMP) and malate dehydrogenase (MDH) generation tasks. Computational experiments confirmed that our method outperformed various deep generative models (e.g., generative adversarial net, variational autoencoder, and GPT model without the proposed training strategy) on these tasks, demonstrating the effectiveness of our multi-likelihood optimization strategy.

## Introduction

Proteins are essential molecular machines, playing vital roles in maintaining various cellular functions, regulating metabolic pathways, and influencing disease development (Hartl, Bracher, and Hayer-Hartl 2011). Traditional protein engineering, aimed at designing industrial and pharmaceutical proteins, has primarily relied on directed evolution. This method involves random mutagenesis coupled with manual selection to identify variants with desired properties (Li et al. 2020). However, the vast protein sequence space and the slow pace of experimental validation make this process time-consuming and expensive. Computational-aided protein design offers an alternative approach by leveraging algorithms to automatically generate, optimize, and prioritize proteins with desired functions (Tran and Hy 2024). This paradigm significantly accelerates protein design and improves the success rate of discovering novel functional biomolecules.

Among various computational approaches, machine learning (ML), particularly deep learning-based protein design, has garnered significant attention due to its improved modeling accuracy, efficiency in navigating the protein sequence space, and high-throughput productivity (Brandes et al. 2022). Protein design using deep generative models can generally be divided into two categories: structure and sequence-based design. Sequence-based approaches directly generate amino acid sequences with desired functions, whereas structure-based approaches first generate functional 3-D protein structures and then reverse-engineer the corresponding amino acid sequences (Notin et al. 2024). While structure-based design might seem more accurate and explainable due to its direct modeling of protein structures, it is hampered by the limited availability of experimentally determined 3D protein structures, restricting the training of reliable deep generative models. One could argue that predicted structures from tools like AlphaFold (McDonald et al. 2023) and ESMFold (Lin et al. 2022) could be used to augment training data for structural protein generation. However, this approach may introduce noise into the training data, potentially compromising the quality of the resulting models, as not all proteins are well-represented by these structural predictors. Additionally, some proteins and peptides—particularly short, flexible, and dynamic ones—are challenging to capture with a single predicted structural conformation, making the structure–function relationships difficult to model (McDonald et al. 2023). Due to these challenges, sequence-based design remains a major method for efficiently generating functional proteins.

Almost all mainstream deep generative model frameworks (i.e., variational autoencoder, generative adversarial net, flow-based, diffusion-based and neural language models) have been adopted for functional protein sequence generation (Harshvardhan et al. 2020). These methods typically involve maximizing the likelihood or its lower bound of training protein sequences within the amino acid sequence space. In other words, the generated protein sequences should closely resemble those in the training data in terms of their ordered amino acid composition. However, optimizing solely within the amino acid space may not be ideal. Here we provide two justifications for this. First, amino acid changes (i.e., mismatches, insertions and deletions) in protein sequences (i.e., “genotypes”) don’t necessarily cause their function changes (i.e., “phenotypes”). For instance, consider a generative model trained on an antibiotic protein *A* (sequence: *KTLKV*) that generates two variants, *A*_1_ (sequence: *KTLKR*) and *A*_2_ (sequence: *KTLKM*). Although both *A*_1_ and *A*_2_ differ from *A* by one amino acid, and thus have the same training loss, *A*_1_ might be identified as an antibiotic while *A*_2_ might not. Penalizing both variants equally during training fails to capture the functional similarity between *A* and *A*_1_. In addition to this “genotype-phenotype” issue, the amino acid sequence space is discrete, and functional protein distributions within this space can be highly multimodal, hindering distribution learning (**Fig.1a-b**). These challenges suggest that we should optimize the likelihood of training data beyond the amino acid sequence space. Recent advances in foundational models, such as GPT- and BERT-style models trained for protein sequences (e.g., ProtGPT, ProGen, ESM series ESM1 and ESM2, Prot-BERT), have revolutionized computational protein modeling (Ferruz and Höcker 2022; Lin et al. 2022). Specifically, the BERT-style ESM model embeds protein sequences into a latent space where proteins with similar functions tend to have similar latent embeddings, effectively serving as an indirect protein function checker (Lin et al. 2022). We hypothesize that the distribution of functional proteins in this latent space is more organized than in the amino acid sequence space, making it more suitable for deep generative modeling (**Fig.1a-b**). Inspired by this, we propose a multi-likelihood space optimization strategy for functional protein generation. Specifically, we trained GPT models by simultaneously optimizing the training data likelihoods in both the original amino acid sequence space and the ESM-derived latent space. As the loss from the latter space is not differentiable, we used policy gradient during training. Our training strategy can also be viewed as a form of knowledge distillation from ESM to GPT, where knowledge transfer is achieved through dynamically reweighting the gradients of generated samples during training. We applied our method to generate antimicrobial peptides (AMPs) and malate dehydrogenases (MDHs), demonstrating that our approach outperformed the original GPT model fine-tuned on functional proteins sets as well as several baseline deep generative models in these tasks. Our training strategy is flexible and can be adopted by other generative models that explicitly model the likelihood of training data. We envision that our approach could serve as a plug-in during deep generative model training to improve the functional protein generation quality.

**Figure 1:**
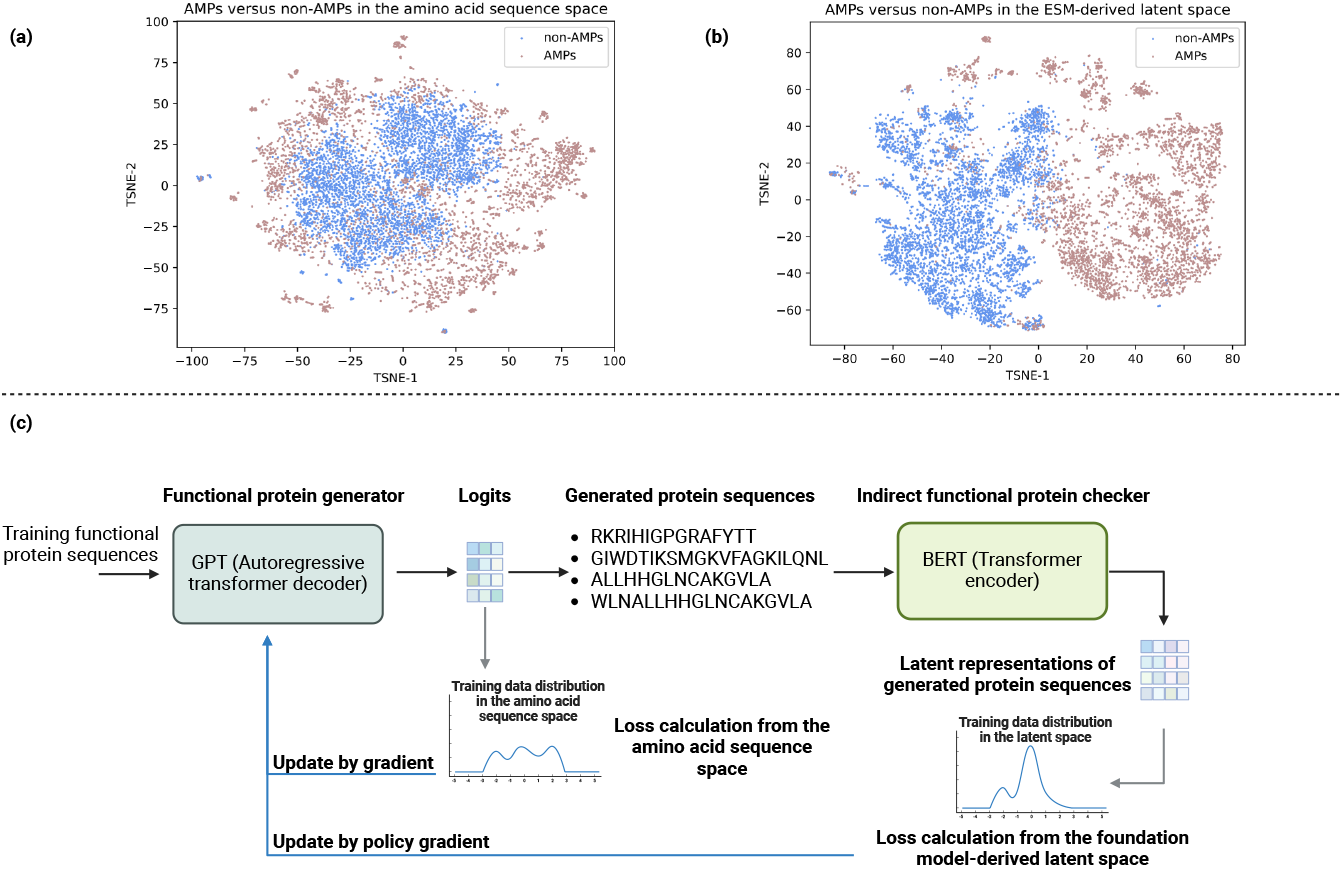
**(a)** A sequence similarity matrix was constructed using 5,000 antimicrobial peptides (AMPs) and 5,000 non-AMPs through local sequence alignment. T-SNE was then applied to convert the similarity matrix into a 2D space. **(b)** For the same set of AMPs and non-AMPs, ESM2 was used to generate their latent representations, which were subsequently reduced to two dimensions using T-SNE. Compared to the 2D space derived from sequence similarity, the 2D latent space induced by the ESM model showed improved separation and clustering of AMPs and non-AMPs. **(c)** Our proposed framework for functional protein sequence generation incorporates an additional protein BERT model into its training process. This allows for the training of an enhanced deep generative model (e.g., a GPT model) by simultaneously optimizing the training data likelihoods from both the original protein sequence space (i.e., the amino acid sequence space) and the BERT-derived latent space.

## Related Work

### Knowledge distillation

Knowledge distillation is a process where a student model (typically smaller and more efficient) is trained to replicate the behavior of a teacher model (typically larger and more complex). By converting the outputs of the teacher model into “soft labels” for the student model to learn, the student can effectively learn and internalize the knowledge from the teacher. This technique is often employed for model pruning, transfer learning, and ensemble learning. For a more comprehensive review of knowledge distillation, please refer to this review paper (Gou et al. 2021).

### Deep generative models for functional protein design

To advance protein design, various deep learning methods have been developed based on deep generative model frameworks such as Generative Adversarial Nets (GANs), Variational Autoencoders (VAEs), neural language models, flow-based models, and diffusion models. Representative structure-based protein design approaches include conditional VAEs (Schmitt et al. 2022), guided diffusion (Gruver et al. 2024), Regression Transformer, and ProteinNPT (Notin et al. 2023). Conversely, sequence-based protein design often benefits from lower modeling complexity and better availability of labeled data for training. For antimicrobial peptide (AMP) generation, methods have been developed using GANs (e.g., AMPGAN, Feedback-GAN), VAEs (e.g., PepCVAE, HydrAMP) (Szymczak et al. 2023), GPT-based models, and hybrid approaches (e.g., GAN + diffusion). Additionally, GAN (ProteinGAN) (Repecka et al. 2021) and GPT architectures have been applied to malate dehydrogenase (MDH) design and the discovery of novel MDH variants.

### Foundation models for protein sequence modeling

Pre-trained large language models, such as ProtGPT (Ferruz, Schmidt, and Höcker 2022), ProGen (Ferruz and Höcker 2022), ESM (ESM1 and ESM2) (Lin et al. 2022), and ProteinBERT (Brandes et al. 2022), have been trained on massive protein sequence datasets in a self-supervised manner to perform general protein sequence generation and protein sequence representation learning. Although these models are trained for general purposes, they can be easily fine-tuned on smaller functional protein sets to perform specific down-stream tasks, such as generation and property prediction.

## Methodology

We first provide an overview of the general workflow of our method, which comprises a deep learning-based generator for protein sequence generation and a pretrained foundation model for protein sequence representation learning (**Fig.1c**). The generator can be any deep generative model framework capable of explicitly modeling the training data likelihood or its lower bound (e.g., variational autoencoder, flow-based models, diffusion-based models, or neural language models). While the typical approach to training a generator for protein sequence generation involves maximizing the likelihood of training amino acid sequences, the inclusion of a pre-trained protein foundation model allows us to derive an alternative training data distribution—specifically, the distribution of latent representations of the training proteins—thereby providing an additional optimization target for the generator. For simplicity and without loss of generality, we assume our generator is a GPT model (i.e., autoregressive transformer decoder) and the pre-trained protein foundation model is a BERT model (i.e., transformer encoder) (Vaswani 2017). Below, we introduce some basic concepts and notations related to GPT and BERT models before detailing our generative model training strategy.

### GPT for functional protein sequence generation

Given a functional protein set *D* = {*X*_1_, *X*_2_, …, *X*_*n*_} containing *n* sequences for a GPT model to train, the model takes each sequence *X*_*i*_ as its input and utilizes a transformer decoder (with causal mask) to autoregressively perform the next-token probability prediction *p*(*X*_*i,j*_ |*X*_*i,j* −1_, *X*_*i,j*− 2_, …, *X*_*i*,2_, *X*_*i*,1_). Here, *X*_*i,j*_ stands for the *j*th token in sequence *X*_*i*_, and the probability of generating token *X*_*i,j*_ in *j*th position is conditioned on the previous tokens *X*_*i,j* −1_, *X*_*i,j*− 2_,…, *X*_*i*,2_, and *X*_*i*,1_. The standard loss function to be minimized for training such a model is the negative log likelihood (NLL) of the training data in the token (i.e., amino acid) space, which can be written as:

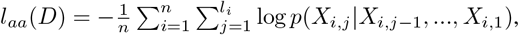

where *l*_*i*_ is the amino acid sequence length for protein *X*_*i*_. As NLL is equivalent to the cross-entropy loss, *l*_*aa*_(*D*) practically is calculated by the cross entropy between the input sequences and the generated sequences by the GPT model. Suppose we input the training protein sequences to the GPT model and obtaining the generated sequences 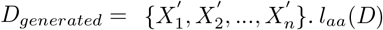 can be also written as:

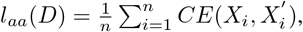

where 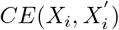 is the cross-entropy loss between two sequences *X*_*i*_ and 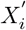.

### BERT for protein representation learning

The BERT model learns latent embeddings for protein sequences by processing masked protein sequences using a transformer encoder and predicting the correct amino acids at the masked positions. Given a pre-trained protein BERT model *F* and a protein sequence *X*_*i*_, *X*_*i*_’s first token representation in the last transformer layer of *F* is defined as the protein representation of *X*_*i*_. It should be noted that there are other ways (e.g., mean of the latent representations of all tokens in the last transformer layer) to derive a protein representation, we only used the first token representation here for simplicity. For clarity, we use *F* (*X*_*i*_) ∈ ℝ^*m*^ to denote this protein representation, where *m* stands for the dimension of this latent vectorized representation. By generating representations for all functional proteins from D, we derive an alternative empirical distribution *D*_*latent*_ = {*F* (*X*_1_), *F* (*X*_2_), …, *F* (*X*_*n*_)}for the GPT model to learn from.

### Proposed training scheme

Given the functional protein sequence set D as well as its corresponding latent representation set *D*_*latent*_, our training strategy for the GPT model involves the maximum likelihood estimation from the empirical distributions of *D* (in the amino acid sequence space) and *D*_*latent*_ (in a BERT model-derived latent space). Maximum likelihood estimation for the amino acid sequence space can be achieved via minimizing *l*_*aa*_(*D*) described above. As *l*_*aa*_(*D*) is differentiable with respect to the learnable parameters in the GPT model, this minimization can be achieved by using (mini-batch) gradient descent to update the GPT model.

We now move to describe the way to perform maximum likelihood estimation for *D*_*latent*_. The process starts from inputting the training protein sequences to the GPT model and obtaining the generated sequences 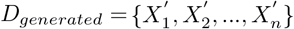. We then pass *D*_*generated*_ to the pre-trained protein BERT *F* to get the latent representations 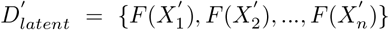 for these generated sequences. Note that the BERT model is pretrained and won’t be updated during our optimization process. As *F* (·) induces a continuous vector space, maximizing the likelihood given *D*_*latent*_ corresponds to minimizing the mean squared error between *D*_*latent*_ and 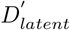 :

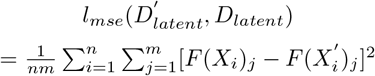

where *F* (*X*_*i*_)_*j*_ stands for the *j*th element of *F* (*X*_*i*_).

Putting the two minimization losses together, we have the final optimization loss for the GPT mode:

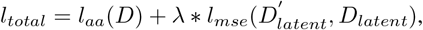

where *λ* is a hyperparameter to balance two losses. Ideally, given the learnable parameters *w* of the GPT model, we want to calculate the gradient of *l*_*total*_ with respect to *w* to perform gradient descent. However, 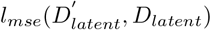 is not differentiable with respect to *w*. To address this, we use policy gradient (Sutton and Barto 2018) to replace the “invalid” gradient of *l*_*mse*_ to *w*. Specifically, in the context of reinforcement learning, we formulate the GPT as a policy network and the BERT model as the environment the policy network interacts with. We define the reward as the negative 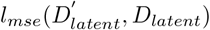 (i.e., we want to minimize the reward here). Then, sample-wisely, the policy gradient of the reward from the generated sample 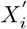 with respect to *w* is the multiplication between the reward and the log likelihood of 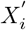 from the GPT model (i.e., the negative cross-entropy between 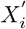 and *X*_*i*_):

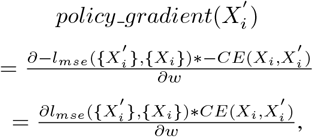

where ∂ is the partial derivative symbol. Now the gradient of *l*_*total*_ with respect to *w* for the generated sample 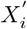 can be written as:

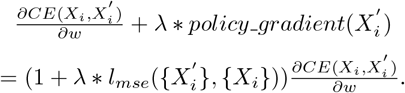

We then can use this gradient to update the parameters w of the GPT model. By looking at the gradient form, we can see that 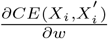 is just the original gradient for training the GPT model. Our proposed training scheme extends it by adding a dynamic reweighting strategy (i.e., by the weighting term 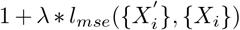 during each iteration of the GPT model training. This can be considered as a form of knowledge distillation from the BERT model to the GPT model. Generated sequence 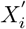 having a larger distance to its training template *X*_*i*_ in the BERT-derived latent space is considered to be a more severe error and will receive a bigger attention (i.e., larger gradient) during the training. We believe that by penalizing functionally different (determined by the BERT model) sequence generations more during the GPT model training, our generator can achieve an improved functional protein generation quality.

### Training details

We utilized the ProGen2-large model, a 2.7 billion-parameter GPT model pre-trained on protein sequences from UniProt, as the initialization for our functional protein generator. For deriving the latent space distribution of proteins that the GPT model aims to capture, we selected the ESM2 with 33 layers as the BERT model. For both the AMP and MDH generation tasks, we trained the GPT model using the AdamW optimizer with betas of 0.9 and 0.999, and an epsilon value of 1e-8. The AMP generation task was trained for 50 epochs with a batch size of 16 and a learning rate of 1e-5, while the MDH generation task was trained for 30 epochs with a batch size of 4 and a learning rate of 1e-5. During training, a warm-up schedule was implemented, the learning rate is warmed up over 5% of the total training steps to a peak value of 1e-5, followed by a gradual decrease. Before inputting the protein sequences into the GPT and ESM2 models, we added a start token and an end token to the beginning and end of each sequence, respectively. To augment the number of training samples for the GPT, we also used the reversed protein sequences, training the GPT model to generate both the standard sequences (from N-terminus to C-terminus) and their flipped versions (from C-terminus to N-terminus). For the policy gradient, we set *λ* ∈ {1, 10, 100}. In addition to the original reward defined in the subsection **Proposed training scheme**, we also introduced a reward variant, defined as the original reward subtracted by a baseline. The baseline was calculated as the exponential moving average (EMA,with its hyperparmeter set to 0.9) of the historical original reward during GPT training. We randomly split the training data, setting aside a percentage as a test set. During the GPT model training, we evaluated the perplexity of the test set, and the model with the best test data perplexity was saved for further evaluation.

## Experiments

To evaluate our method, we selected two distinct types of proteins for benchmarking: antimicrobial peptides (AMPs) and malate dehydrogenases (MDHs). AMPs are short chains of 1-100 amino acids that can exhibit antibiotic activity, offering a potential strategy to kill bacteria and treat infections (Wan et al. 2024b; Wong, de la Fuente-Nunez, and Collins 2023). MDHs are key enzymes in the central oxidative pathway and the tricarboxylic acid cycle, catalyzing the conversion of malate to oxaloacetate using NAD+ as a cofactor. These two protein types were chosen due to their significant relevance to human health: AMPs present a promising approach to combat antibiotic resistance, while MDHs play a crucial role in cellular metabolism.

### Experimental setups

#### Datasets

For the AMP generation case, we collected experimentally verified AMP sequences from five AMP databases (i.e., ADP3 (Wang, Li, and Wang 2016), DRAMP (Shi et al. 2022), LAMP2 (Ye et al. 2020), DBAASP (Pirtskhalava et al. 2021) and dbAMP (Jhong et al. 2022)), resulting in a total of 42,210 AMP sequences for training. To draw **Fig.1**, we randomly sampled 5,000 sequences from the curated AMPs and 5,000 non-AMPs from Ma et al. (Ma et al. 2022). For the MDH generation case, we obtained the MDH data from Zeng et al. (Zeng et al. 2023) study, which comprises 16,706 MDH training sequences and 214 MDH test sequences. Additionally, to construct the MDH predictor to evaluate the performance of the generative model, we obtained another dataset from the same study, consisting of a balanced set of MDH and non-MDH sequences (*n* = 16, 706 + 16, 706). From this dataset, we randomly sampled 4,500 sequences (13.47%) for training the MDH predictor, 500 sequences (1.50%) for validation, and used the remaining 28,412 sequences (85.03%) for testing.

#### Metrics

We used three metrics to evaluate the generation quality of various deep generative models: **(1) Prediction performance from external protein function predictors**. The primary goal of functional protein design is to generate biomolecules with desired properties. Therefore, better generation models should yield superior results when evaluated using relevant protein function predictors. For AMP prediction, we employed two models: Macrel (Santos-Júnior et al. 2024), an AMP classification model, and APEX (Wan et al. 2024a), an antimicrobial activity regressor model. In Macrel, a higher predicted probability indicates that the corresponding input sequence is more likely to be antimicrobial. APEX, on the other hand, outputs multiple minimum inhibitory concentrations (MICs, unit: *µmol/L*) against several bacteria for a given sequence, with the median MIC representing the overall antimicrobial activity. Unlike the AMP classifier, a lower MIC value indicates higher antimicrobial activity. We selected Macrel and APEX because they have been validated for practical AMP virtual screening, and their reliability has been confirmed through experimental verification. Using Macrel, we calculated two metrics: Avg macrel and P marcel. Avg macrel represents the mean predicted probability for a set of protein sequences, while P macrel denotes the percentage of proteins with a predicted probability greater than 0.5 within the set. Similarly, we derived two metrics from APEX: Avg apex, which calculates the mean of median MICs for the sequences, and P apex, which indicates the percentage of proteins with a median MIC of less than 80 *µmol/L* in the set. For MDH prediction, we built a predictor using a BERT model fine-tuned on the 33-layer ESM2 model. This model was trained for 10 epochs using a learning rate of 1e-6, a batch size of 16, and the AdamW optimizer. The best-performing model on the test set was saved for MDH prediction. We used the same definitions for Avg macrel and P macrel to derive Avg mdh and P mdh for evaluating MDH generation quality. **(2) Sequence similarity analysis**. To assess the sequence similarity between generated and training protein sequences, we compared a generated protein set *B* with a training protein set *A*. The sequence similarity of a protein sequence *i* in B with respect to *A* is defined as the maximum normalized Smith-Waterman alignment score between sequence *i* and any protein sequence *j* in *A*. The Smith-Waterman algorithm is a local sequence alignment approach used to compare the similarity of two DNA or protein sequences. Detailed information on the normalized Smith-Waterman alignment score can be found in the APEX paper. For the generated protein set *B*, we calculated the sequence similarity for all sequences from *B* and reported the mean and standard deviation of these similarities. Defining an appropriate range of sequence similarity is not trivial. A high similarity score (e.g., mean score *>* 0.9) indicates that the generator captures the training distribution well, but it may also suggest that the generator is not exploring a sufficiently large protein space, limiting the potential for discovering novel sequences. Conversely, a low similarity score (e.g., mean score *<* 0.3) suggests that the generator fails to learn the empirical distribution of the training data. In previous studies, a sequence similarity of less than 75% was used to define novel protein sequences (Wan et al. 2024a). We followed this empirical cutoff to establish a sequence similarity range of 60% to 80%, balancing sequence space exploration and exploitation. An effective sequence generation should fall within this defined range. **(3) MSA Entropy**. To assess the diversity within the generated protein sets, we performed multiple sequence alignments using MAFFT version 7 (https://mafft.cbrc.jp/alignment/server/). We then calculated the column-wise entropies of the MSA results using ProDy (http://www.bahargroup.org/prody/). MSA entropy was defined as the summation of these column-wise entropies. Higher MSA entropy indicates greater diversity within the generated protein set.

#### Baseline methods

For the AMP generation task, we selected five models for comparison: AMPGAN (GAN-based), VAE-basic (VAE-based), PepCVAE and HydrAMP (conditional VAEs), and Diff-AMP (GAN+diffusion). For the MDH task, ProtGPT was fine-tuned on MDH data for comparation. The MDH dataset (n=16,706) was clustered by using CD-HIT and four clusters as cluster765, cluster2029, cluster5987, and cluster7477, with the number inside the name indicating the number of sequences in that cluster (e.g., cluster765 contain 765 sequences) were obtained. These four clusters were separately used to fine-tune ProtGPT to obtain four MDH generation models. We simply named them GTP-cluster765, GTP-cluster2029, GTP-cluster5987 and GTP-cluster7477. ProGPT without fine-tuning was also used as a baseline for comparison. In addition, we also incorporated a Random generator that used the amino acid composition (AAC) information of training MDH sequences and randomly generated sequences by sampling amino acids from the training AAC distribution.

#### Our method

For our GPT model, we employed three different sequence generation strategies: (1) generating from N-terminus to C-terminus, (2) generating from C-terminus to N-terminus (i.e., reverse generation), and (3) an even mixture of both (1) and (2). For AMP generation, we named our model AMPGPT C terminal, AMPGPT N terminal and AMPGPT Mixture, respectively. Note that in the method section, we also derived a reward by subtracting a baseline. We named GPT models trained by this variant reward by AMPGPT C terminal EMA, AMPGPT N terminal EMA and AMPGPT Mixture EMA. As an ablation study, we compared our GPT models to those trained without the policy gradient. These models were named as BaseGPT C terminal, BaseGPT N terminal and BaseGPT Mixture. Similarly, for the MDH generation, we used the same naming strategy and replaced the “AMP” by “MDH”. For the AMP task, we counted the frequencies of dipeptides formed by the first two amino acids starting from the N-terminus and C-terminus of the training set sequences, respectively. We then sampled dipeptides according to these frequencies and used them as input to generate AMP sequences. Specifically, when generating from the N-terminus, we sampled dipeptides based on the N-terminal dipeptide frequency, and when generating from the C-terminus, we sampled dipeptides based on the C-terminal dipeptide frequency. For the MDH task, the same strategy was used except the decapeptide frequency was used instead of dipeptides. For all generation methods, we generated 10,000 AMPs and 4,000 MDHs for comparison.

### AMP generation evaluation

To demonstrate the effectiveness of using the policy gradient to minimize the latent space distance between training and generated sequences, we visualized the training process. That is, with the increase of the training epochs, the generated AMP sequences became closer to their training templates in the ESM2 space (**Fig. S1**). After confirming the effectiveness of the policy gradient optimization, we evaluated the AMP generation quality for different deep generative methods. As our model involves the hyperparameter *λ* for weighting the policy gradient during training. We used the AMP classification model Macrel to evaluate our GPT models under different *λ*s and selected the best GPT models according to Avg macrel. After determining the *λ*, we first studied the AMP generation quality by evaluating the AMP prediction results of the generated sequences by Macrel and APEX. As an ablation study, we first compared the AMP predictions for the generations from AMPGPTs (proposed method), AMPGPT EMAs (proposed method with a baseline subtraction from the reward), and BaseGPTs (no using policy gradient). For the prediction under Macrel, we observed that the predictions for the generations from BaseGPTs, AMPGPTs and AMPGPT EMAs are relatively comparable (**Table1, Figs. 2 and S2**). Even if on some cases the BaseGPTs outperformed AMPGPTs and AMPGPT EMAs, statistical tests showed that the differences were not significant. However, when evaluating the AMP prediction by APEX, we observed that AMPGPTs and AMPGPT EMAs significantly outperformed the BaseGPTs (**Table1, Figs. 2 and S2**), demonstrating the effectiveness of our proposed training strategy. Between AMPGPTs and AMPGPT EMAs we did not observe a consistent winner. When comparing AMPGPTs and AMPGPT EMAs to other baseline deep generative models, we found that our methods consistently achieved significantly better predictions from APEX and Macrel. This suggest that our method has a better capability of generating antibiotic-like sequences.

**Table 1:**
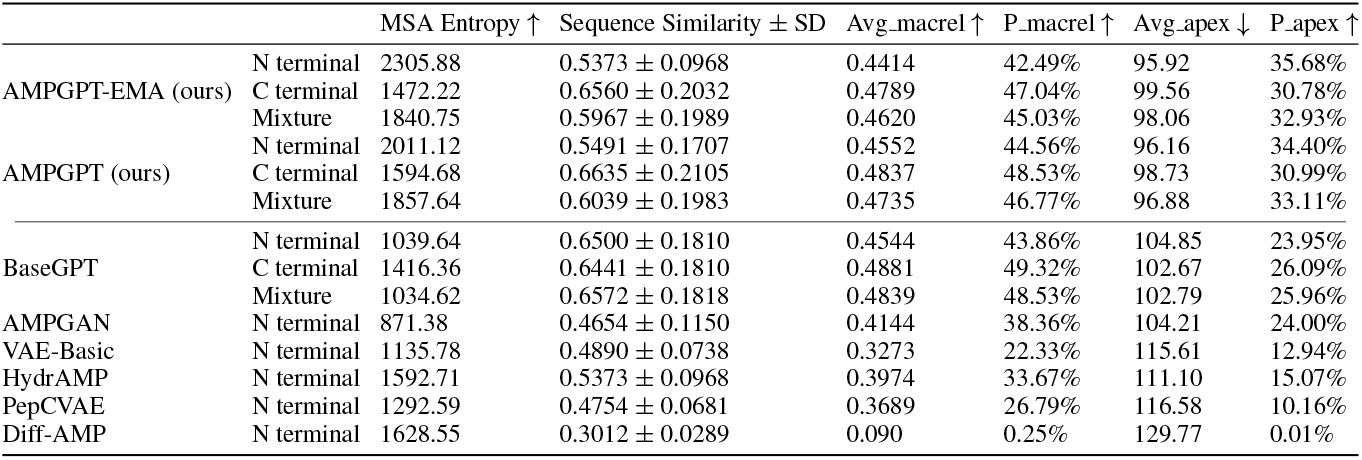
A summarization of metrics for evaluating generated AMPs from different methods. SD stands for standard deviation. Arrow directions indicate the improvement directions for the corresponding metrics.

**Figure 2:**
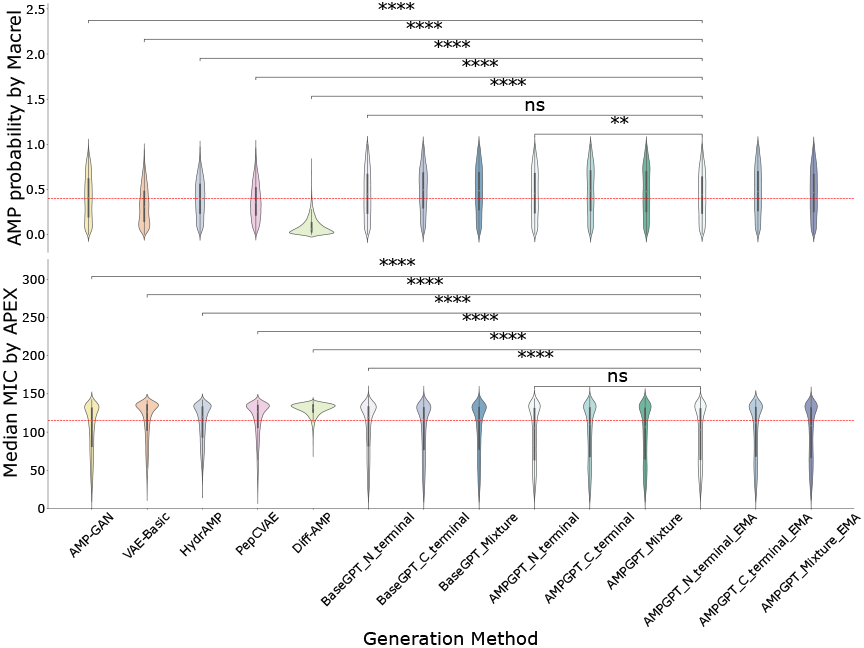
Violin plots for comparing AMP probability prediction distribution and MIC prediction distribution for different AMP generation methods. AMPs should have high AMP probability prediction and low MIC prediction. Statistical tests were used to compare the distribution between our the AMPGPT N terminal EMA model and the corresponding baseline models. The statistical significance of the results was evaluated using the Mann-Whitney U test. ns: Not significant (5.00e-02 *<* p ≤ 1.00e+00); *: Significant (1.00e-02 *<* p ≤5.00e-02); **: Highly significant (1.00e-03 *<* p ≤ 1.00e-02); ***: Very highly significant (1.00e-04 *<* p ≤ 1.00e-03); ****: Extremely significant (p ≤ 1.00e-04). The red line for the upper figure is *y* = 0.4 and the red line for the lower figure is *y* = 115. We drew them to facilitate the comparison among different methods.

Furthermore, a deeper analysis of the generated AMP sequences from AMPGPT and AMPGPT EMA variants revealed that they exhibited a sequence similarity ranging from 0.5 to 0.67 (**Table1**). Since we defined that a sequence similarity ranging from 60% to 80% reflects a good sequence space exploration and exploitation. We can see that compared to the sequence similarities resulted by other deep generative models, the sequence similarities derived from our methods overlapped better to this range. This suggests that our method is more capable of capturing the underlying distribution of the training data, while also generating novel yet functional sequences. In addition, we observed that the sequence similarities of AMPGPT EMA and AMPGPT variants tend to be lower than those of BaseGPT variants. This aligns with our anticipation as AMPGPT EMAs and AMPGPTs were trained to maximize the likelihood of training data not only from the amin acid sequence space but also from the ESM2 space, and hence AMPGPT EMA and AMPGPT generations were supposed to be sequentially more different from training ones than those generated from BaseGPTs. When using MSA entropy to quantify the sequence diversity within a generation sequence set, we also observed that AMPGPTs and AMPGPT EMAs generally outperformed others. To further visualize the distribution of generated AMP sequences and training AMPs, we employed t-SNE analysis to visualize them in the ESM2 latent space. The results, presented in **Fig. S3** demonstrated that the generated AMPs effectively covered the training data well. Taken together, we showed that our multi-likelihood optimization strategy for the GPT model led to improved and diverse generation of AMP sequences.

### MDH generation evaluation

After demonstrating the success of our method applied on short AMP sequence generation, we applied our method to design long MDH protein sequences. By using a BERT-based MDH function predictor, we saw that our MDHGPT and MDHGPT-EMA variants achieved better MDH predictions (i.e., better overall Avg mdh and P mdh) than those from the GPT model without our proposed training scheme and the other GPT models developed in (Nijkamp et al. 2023) (**Table2** and **Figs. S4 and S5**). MDHGPT and MD-HGPT EMA generations also enjoyed a decent sequence similarity (overall ranging from 0.6 to 0.8). While the baseline generation models tended to have better MSA entropies, we argued that these results were more likely to be resulted by generating sequences that were less likely to be MDHs (as reflected by their Avg mdh and P mdh metrics). Taken together, we can see that our proposed method achieved the best MDH generation quality.

**Table 2:**
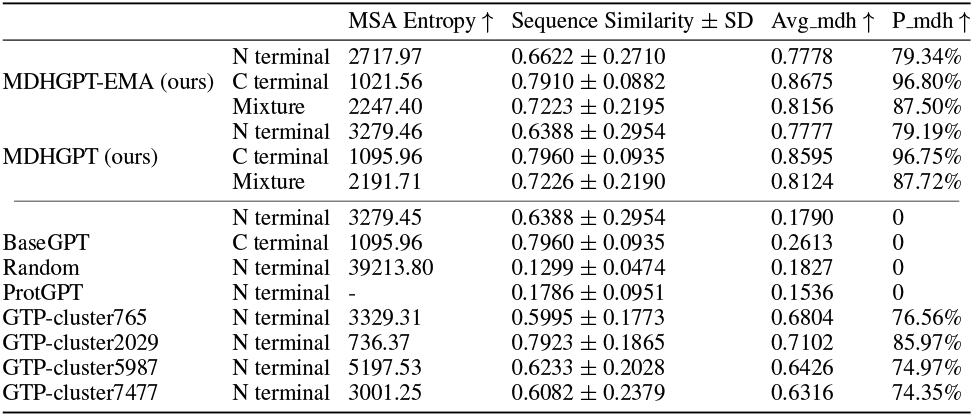
A summary of metrics for evaluating generated MDHs from various methods. SD stands for standard deviation, and the arrow directions indicate the desired improvement for each corresponding metric. N.A. stands for not applicable, as the method generated nonstandard amino acids, which preclude the calculation of MSA.

## Conclusion and future work

In this work, we demonstrated that our multi-likelihood optimization strategy for training deep generative models effectively improves the quality of functional protein sequence generation. Our future direction involves the collaboration with biology researchers to conduct wet-lab validation of our generated sequences.

## Acknowledgements

The corresponding author is a recipient of the Langer Prize by the AIChE Foundation, and acknowledges funding from the IADR Innovation in Oral Care Award, the Procter & Gamble Company, United Therapeutics, a BBRF Young Investigator Grant, the Nemirovsky Prize, Penn Health-Tech Accelerator Award, the National Institute of General Medical Sciences of the National Institutes of Health under award number R35GM138201, and the Defense Threat Reduction Agency (DTRA; HDTRA1-22-10031, HDTRA1-21-1-0014, and HDTRA1-23-1-0001).

