## Supplemental Information for "Improving functional protein generation via foundation model-derived latent space likelihood optimization"

Anonymous Submission

Anonymous affiliation

**Hardware and Software Requirements:**

The experiments in this paper were conducted on a device equipped with two A100 GPUs, each with 80GB of memory. For software requirements, please refer to the resource code instruction, which provides a document with detailed instructions for installing the environment necessary for training our models.

**The hyperparameter**  $\boldsymbol{\lambda}$ **setting:**

$\lambda$ controls the effective of the policy gradient, so we set $\lambda$ ={1,10,100}, and then using predictor to select the best $\lambda$ for models. For both our generation tasks, best $\lambda$ was 10.

**Hyperparameter Settings:**

For GPT generation, there are two hyperparameters temperature *t* and Top *p* to control the quality of generated sequences for each task. For the AMP task, we set *t*=0.8 and *p*=0.95 for AMPGPT and AMPGPT_EMA, while *t*=1 and *p*=0.95 for BaseGPT to mitigate the high duplication rate of sequences. For the MDH task, we set *t*=0.95 and *p*=0.6 for MDHGPT, MDHGPT_EMA, and BaseGPT, as MDH sequences tend to be longer than AMP sequences.

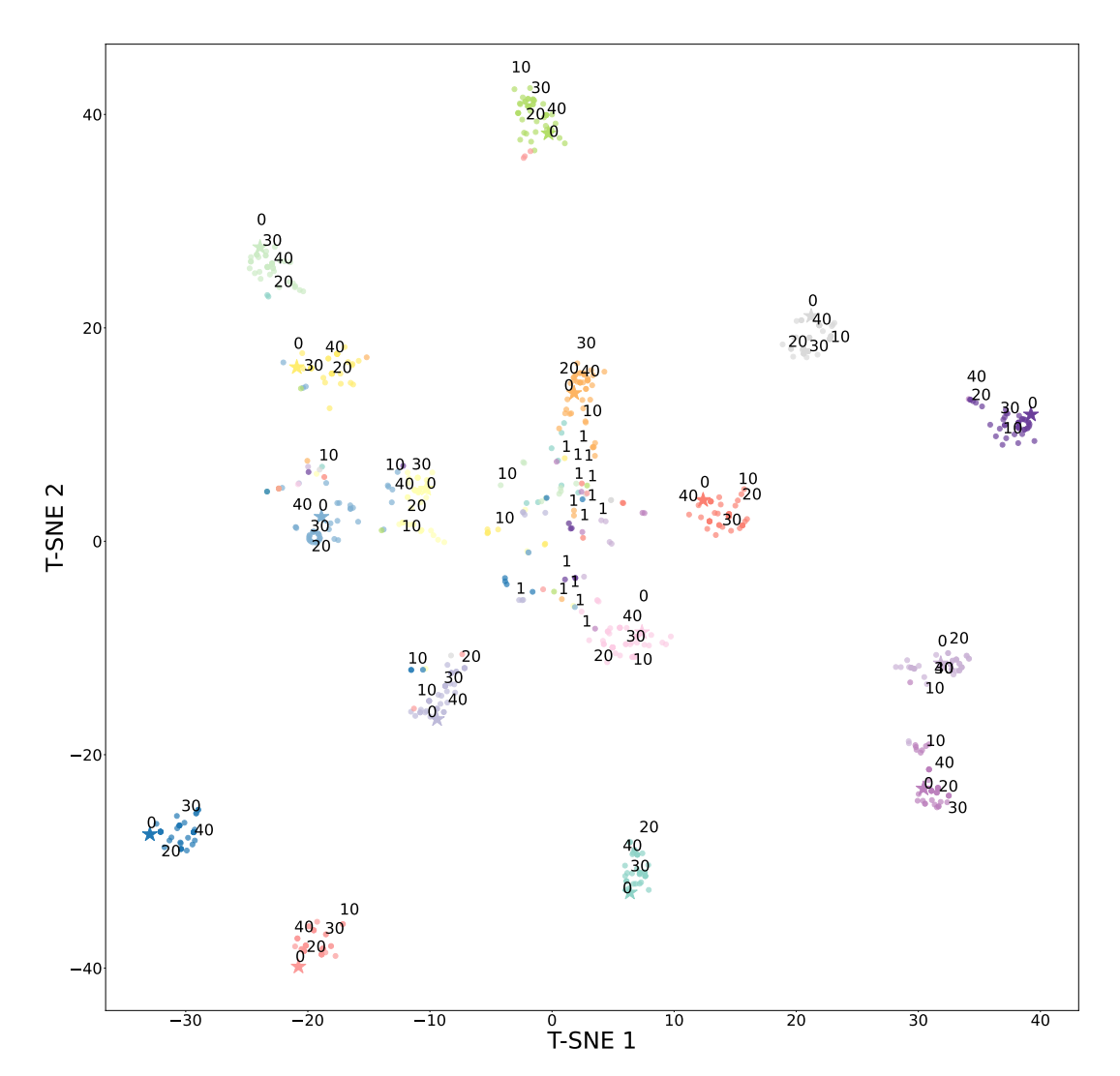

**Fig. S1. Visualization of the generated protein sequences in the ESM2 space during our training process.** Each start point (labeled as 0) represents a training protein sequence, and the round points having the same color as the start point represent the corresponding generated sequences by our GPT model. All sequences were inputted to ESM2. We retrieved the latent representations of all sequences from ESM2 and used T-SNE to plot them in the 2D space. Generated sequences in training epoch 1, 10, 20, 30, 40 were highlighted by texts in the figure. As training progresses, generated sequences became closer to the training ones in the ESM2 space.

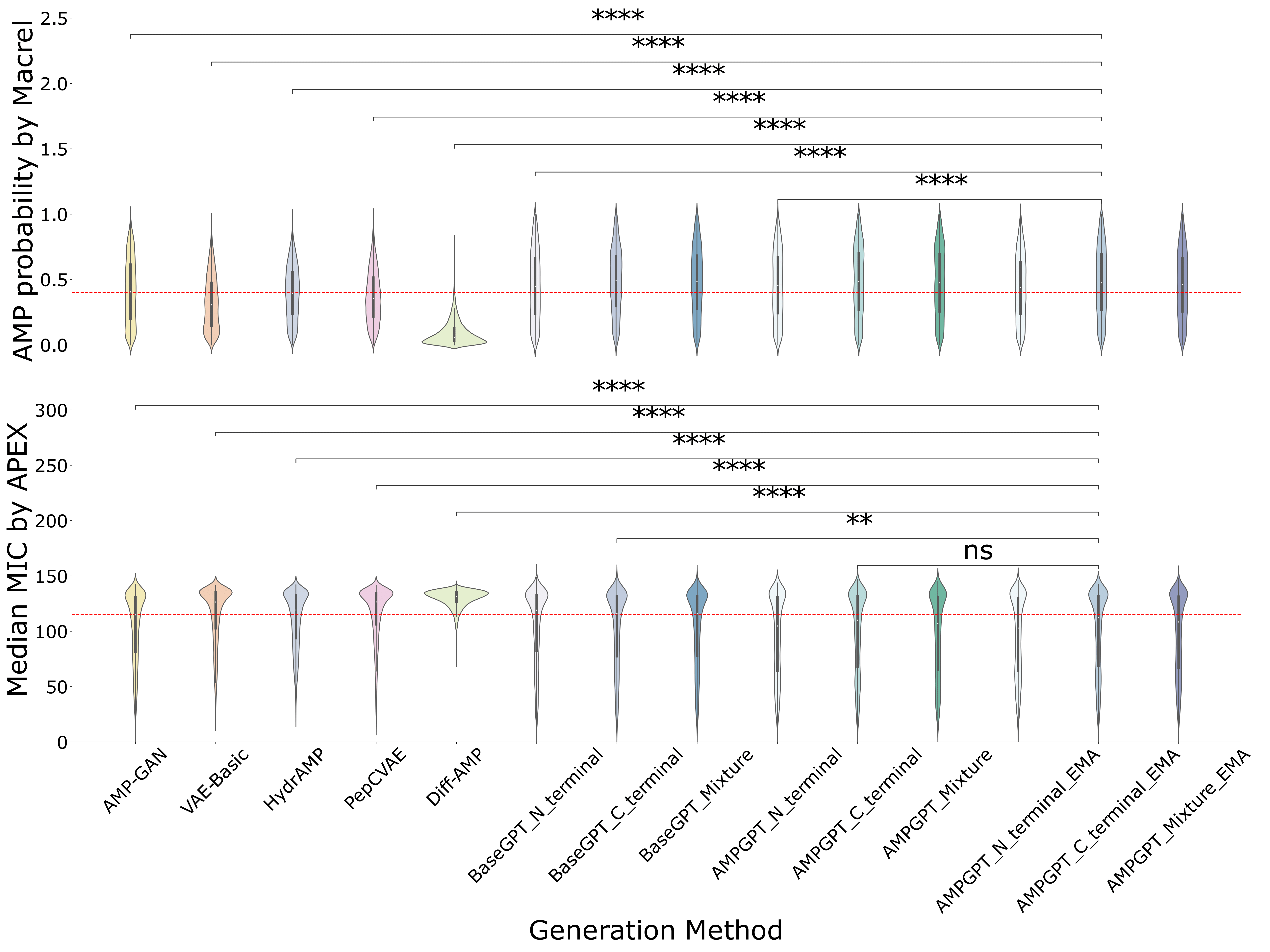

**Fig. S2.** **Violin plots for comparing AMP prediction distribution for different AMP generation methods.** AMPs should have high AMP probability prediction and low MIC prediction. Statistical tests were used to compare the distribution between our the AMPGPT_C_terminal_EMA model and the corresponding baseline models. The statistical significance of the results was evaluated using the Mann-Whitney U test. ns: Not significant (5.00e-02 < p ≤ 1.00e+00); *: Significant (1.00e-02 < p ≤ 5.00e-02); **: Highly significant (1.00e-03 < p ≤ 1.00e-02); ***: Very highly significant (1.00e-04 < p ≤ 1.00e-03); ****: Extremely significant (p ≤ 1.00e-04). The red line for the upper figure is y=0.4 and the red line for the lower figure is y=115. We drew them to facilitate the comparison among different methods.

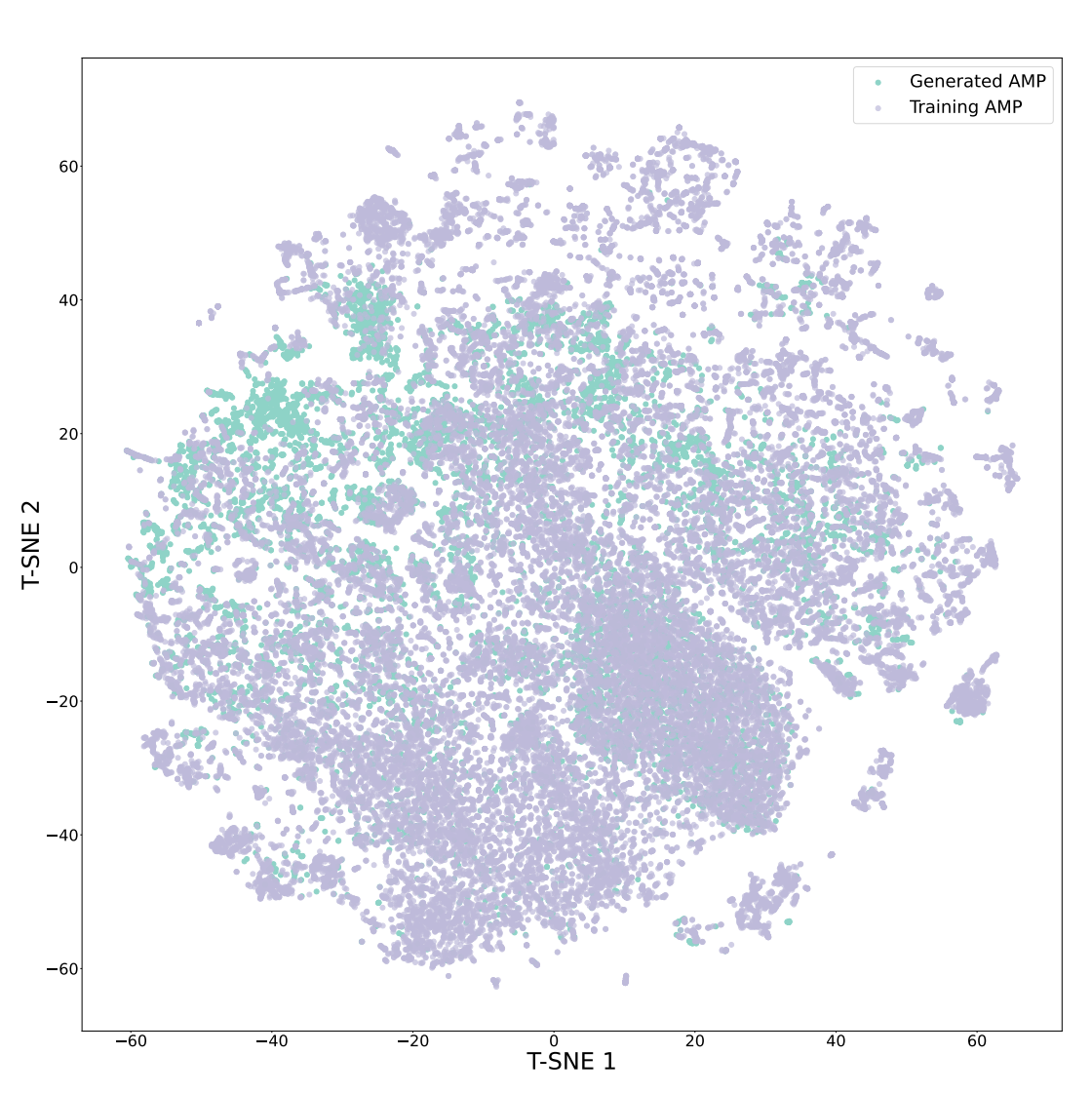

**Fig. S3. T-SNE analysis for the generated AMPs and training AMPs in the ESM2 space.**

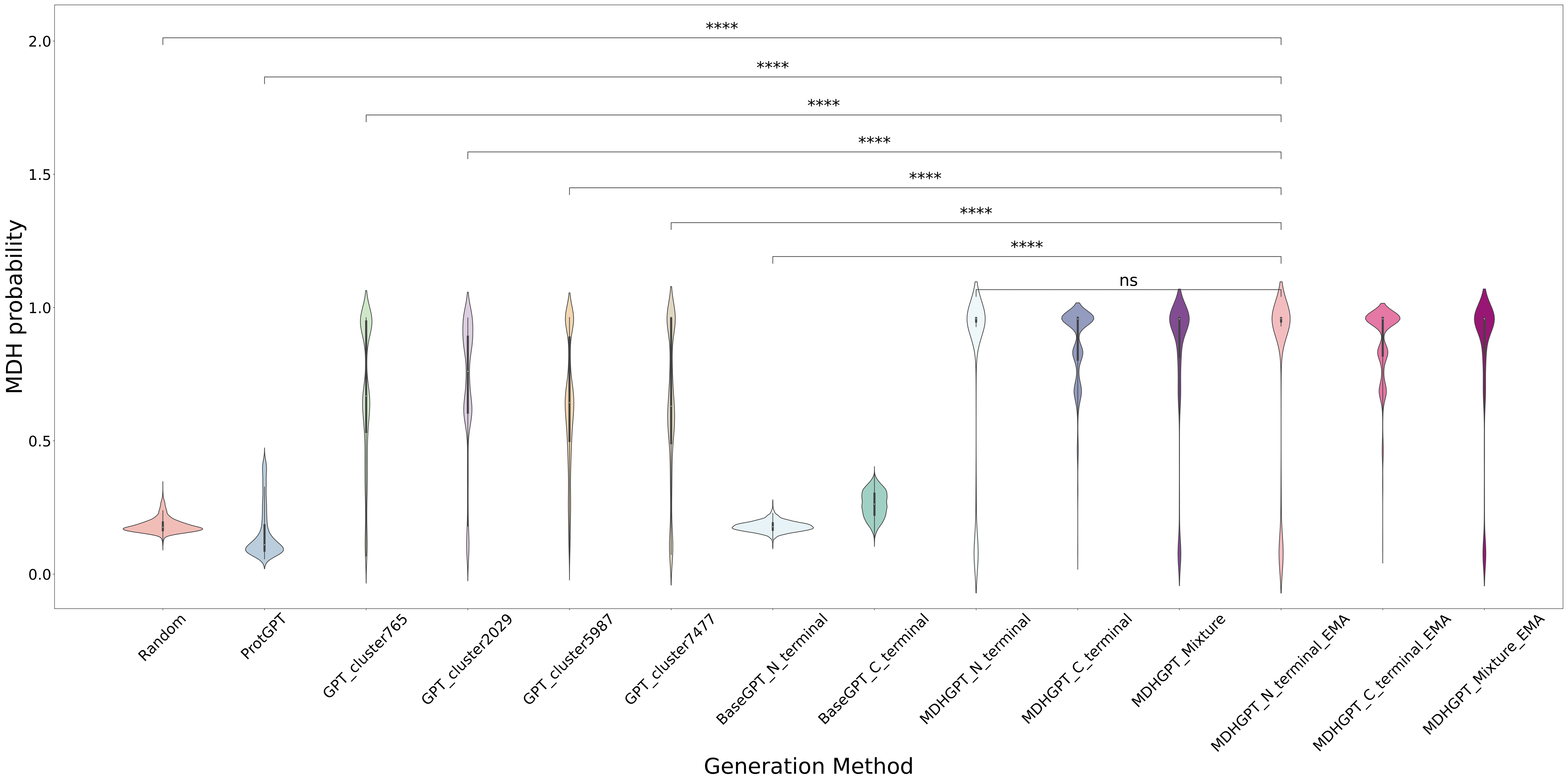

**Fig. S4.** **Violin plots for comparing MDH probability prediction distribution for different MDH generation methods.** MDHs should have high MDH probability prediction. Statistical tests were used to compare the distribution between our the MDHGPT_N_terminal_EMA model and the corresponding baseline models. The statistical significance of the results was evaluated using the Mann-Whitney U test. ns: Not significant (5.00e-02 < p ≤ 1.00e+00); *: Significant (1.00e-02 < p ≤ 5.00e-02); **: Highly significant (1.00e-03 < p ≤ 1.00e-02); ***: Very highly significant (1.00e-04 < p ≤ 1.00e-03); ****: Extremely significant (p ≤ 1.00e-04).

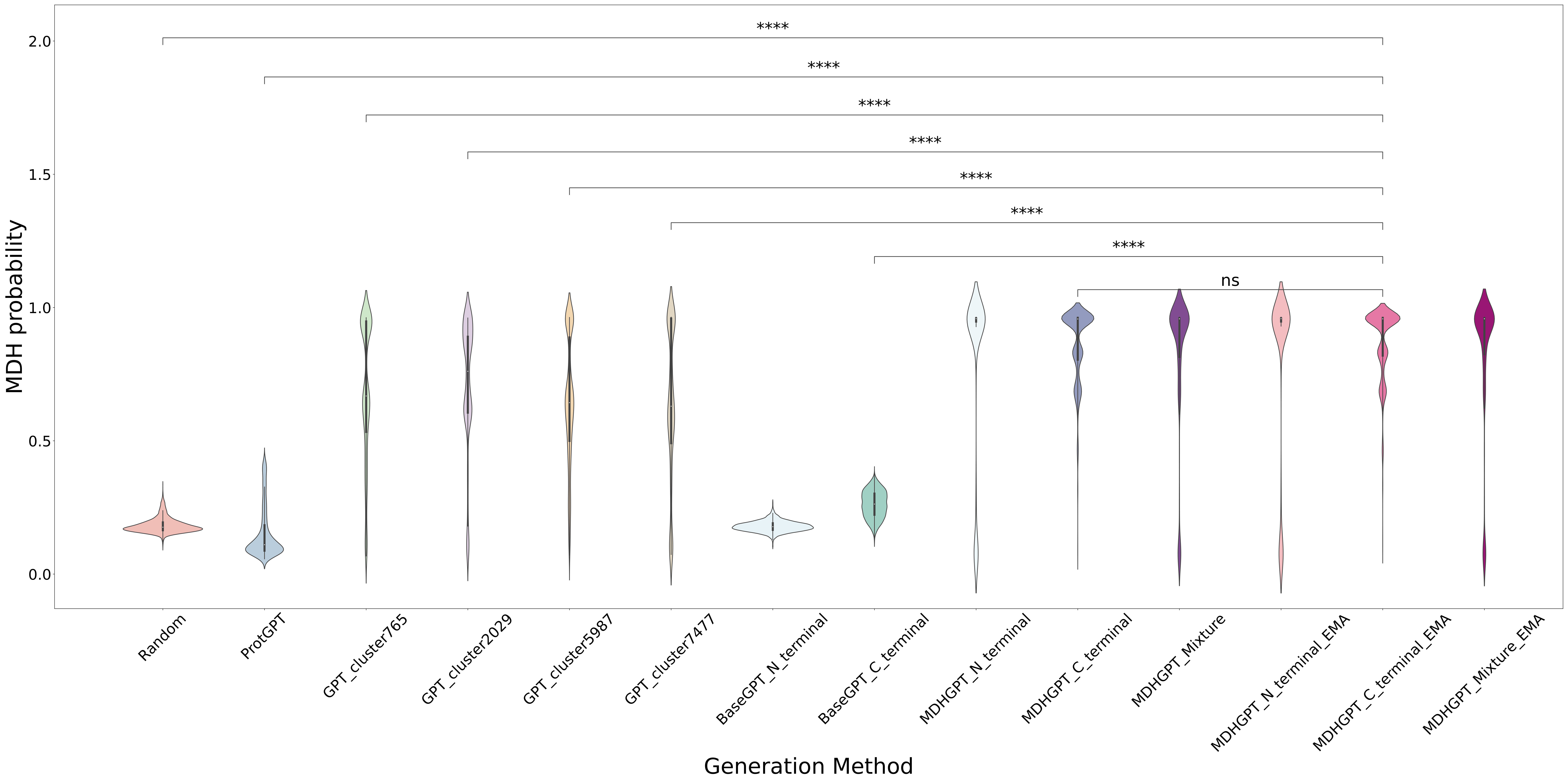

**Fig. S5.** **Violin plots for comparing MDH prediction distribution for different MDH generation methods.** MDHs should have high MDH probability prediction. Statistical tests were used to compare the distribution between our the MDHGPT_C_terminal_EMA model and the corresponding baseline models. The statistical significance of the results was evaluated using the Mann-Whitney U test. ns: Not significant (5.00e-02 < p ≤ 1.00e+00); *: Significant (1.00e-02 < p ≤ 5.00e-02); **: Highly significant (1.00e-03 < p ≤ 1.00e-02); ***: Very highly significant (1.00e-04 < p ≤ 1.00e-03); ****: Extremely significant (p ≤ 1.00e-04).
